# Combining fossil taxa with and without morphological data improves dated phylogenetic analyses

**DOI:** 10.1101/2025.04.07.646048

**Authors:** Mark C. Nikolic, Rachel C.M. Warnock, Melanie J. Hopkins

## Abstract

The fossilized birth-death (FBD) model has become an increasingly popular method for inferring dated phylogenies. This is because it is especially useful for incorporating fossil data into such analyses, integrating fossils along with their age information directly into the tree as tips or sampled ancestors. Two approaches are common for placing fossil taxa in trees: inference based on morphological character data or using taxonomic constraints to control their topological placement. These approaches have historically been treated as alternatives, though combining the two is theoretically possible. For phylogenetic inference of entirely extinct organisms, additional related fossil taxa other than those for which morphology is available are generally overlooked. Here, we implement a combined approach on an empirical dataset for a group of trilobites. We use a morphological matrix and ages for 56 taxa and age information for another 196 taxa from the Paleobiology Database. To evaluate the effects of a combined approach, we conducted FBD dated phylogenetic analyses using a combined approach with morphology and taxonomic constraints and compared them to analyses of taxa with morphology alone. We find that a combined approach yields topologies that are more stratigraphically congruent, substantially more precise parameter estimates (e.g., divergence times), and more informative tree distributions. These findings are a consequence of the substantial increase in stratigraphic age information and a more representative sample of their temporal distributions.

## Introduction

Phylogenetic trees provide necessary context to the study of the evolution of organisms throughout Earth’s history. The fossil and stratigraphic record, in turn, provide the opportunity to directly incorporate the dimension of time in such studies and further inform phylogenies: a quality of this record recognised for at least half a century (Simpson, 1975). Fossils here can directly provide two sources of information: morphology and age. The development of the increasingly popular fossilized birth-death (FBD) process (Gavryushkina et al., 2014; Heath et al., 2014; Stadler, 2010) has provided an especially useful model for incorporating fossil data and some of its idiosyncrasies into phylogenetic analyses (reviewed in Mulvey et al. 2025). The FBD process explicitly considers the lineage diversification (speciation and extinction) process and the fossil sampling/recovery process together, incorporating fossils directly into the tree, while accounting for uncertainty in age and phylogenetic placement in a Bayesian framework.

Fossils can be incorporated as tips into FBD analyses either with or without morphological character data along with their age. When character data are used, the topological placement of fossil taxa can be estimated based on these data with greater precision. Further, combining fossil morphological data with morphological data for extant taxa, along with a molecular alignment, results in the so-called total-evidence analysis (Lee et al., 2009; Pyron, 2010; Ronquist et al., 2012; Ware et al., 2010). Alternatively, fossils without morphological data can be placed in the tree using topological constraints (Barido-Sottani et al., 2023). Usually, these constraints are informed by taxonomy, restricting certain clades to be monophyletic based on higher groupings (e.g., genus or family; (Soul and Friedman, 2015)). Taxonomic constraints thereby allow the age information of fossils to still be appropriately integrated into the analysis in the absence of morphological data. The former strategy has been referred to as the “resolved” FBD model and the latter the “unresolved” FBD model (O’Reilly and Donoghue, 2020). Regardless of the strategy used, the desirable, net-positive effects of fossils (and stratigraphic age information) in phylogenetic analyses is apparent and strongly supported (Barido-Sottani et al., 2020; King, 2021; Mongiardino Koch et al., 2023, 2021; Mongiardino Koch and Parry, 2020).

Morphological data and taxonomic constraints (resolved vs unresolved) have historically been treated as alternative strategies (Barido-Sottani et al., 2023; O’Reilly and Donoghue, 2020), and in practice have been used as such. While the FBD model has been successfully utilised for phylogenetic analysis of entirely extinct groups (Brocklehurst et al., 2022; Ford and Benson, 2020; Martin et al., 2023; Pohle et al., 2022; Wright et al., 2021; Wright, 2017), only the resolved FBD model has been used in this context. The most apparent reason for this is, given the lack of molecular data for fossils, morphological data are needed to infer tree topologies. Fossil taxa are therefore sampled to have the most abundant or diagnostic morphological characters, even if those taxa have a temporal distribution that is appreciably different from the overall distribution of known occurrences for that group. This violates the FBD model assumption of uniform/random sampling and in some cases will provide a biased approximation of the true fossil sampling process (O’Reilly and Donoghue, 2020), which can in turn bias the output of inference using the FBD model.

A combination of the two approaches (morphological data and taxonomic constraints) is theoretically possible. Simulations have indeed shown that a combined approach produces more accurate parameter estimates than either strategy alone (Barido-Sottani et al., 2023). Further, because the distribution of fossil sampling times informs the FBD model parameters – much like other quantitative-paleobiology methods used to infer speciation and extinction rates (like the three-timer (Alroy, 2008) or boundary-crosser rates (Foote, 2000, 1999)) – more fossil data (even as occurrences) are expected to increase precision in divergence times and diversification rate parameters. Occurrence and age information can even influence inferred tree topologies (Barido-Sottani et al., 2020) and appear to hold vital phylogenetic information (Mongiardino Koch et al., 2021). Therefore, using only fossil taxa with abundant morphological characters and ignoring the rest underutilises the information in the fossil record. Combined analyses would therefore allow researchers to take full advantage of the fossil record, particularly for analyses of extinct groups.

Here we implement a combined approach FBD phylogenetic analysis on an empirical dataset for a group of trilobites (morphological matrix and ages) by including stratigraphic age information from occurrences for other members of the group from the Paleobiology Database (PBDB; https://paleobiodb.org). We refer to the combined analysis as a ‘semi-resolved’ analysis and compare it to a ‘resolved’ analysis, where we only include taxa with morphological data coded into the morphological matrix. In doing so, we test the effect of both a substantial increase in stratigraphic age information and a more realistic fossil sampling distribution on FBD analyses in an empirical setting. We explore these effects by conducting a thorough examination of the posterior distributions of inferred divergence times and trees. To compare the quality of topologies when the true tree is unknown, we assess the stratigraphic congruence of trees produced from each analysis. Finally, we explore leaf stability and the quality of consensus trees by assessing how informative of the posterior distribution they are.

## Methods

We conducted both resolved and semi-resolved tip-dated FBD phylogenetic analyses using the morphological character matrix for an order of trilobites from a forthcoming publication of a phylogenetic analysis and revision of the order and is available on Morphobank (http://morphobank.org/permalink/?P5740). This matrix consists of 254 characters for 56 species, spanning ∼125 ma, from the middle Cambrian to the Middle Devonian (the largest number of characters in a trilobite character matrix to date). Age data for the taxa in this morphological matrix were (mostly) obtained as high resolution biozone intervals from the literature and correlated to a global scale based on (Gradstein et al., 2020) to obtain absolute age intervals.

For the semi-resolved analysis, we retrieved all occurrences for every genus in the morphological matrix from the Paleobiology Database (PBDB; https://paleobiodb.org) (Supplementary Material). We cleaned this dataset by removing taxa that were not identified to species and/or were associated with very imprecise stratigraphic intervals (e.g., the entire Cambrian). We also removed species that were in the morphological dataset because we already had higher resolution temporal information from the literature. Following the cleaning procedure, we had a dataset with age information for 194 more species and a combined dataset of 250 species covering the entire range of our lineage of interest with no gaps (Fig. S1). For the PBDB dataset, we then randomly subsampled an occurrence for each species to provide the age interval for that species.

We ran all analyses in the software Beast 2.6.7 (Bouckaert et al., 2019) with the constant rates FBD model as implemented in the Sampled Ancestors (SA) package (Gavryushkina et al., 2014). We ran the ‘resolved’ analyses for 1,000,000,000 MCMC generations and the ‘semi-resolved’ analyses for 2,000,000,000 generations. All analyses had the same priors and parameters, except for the origin time prior. Because of the conflicting hypotheses of the trilobite origin time (Holmes and Budd, 2022; Paterson et al., 2019), we tested the influence of both a uniform and exponential prior on both resolved and semi-resolved analyses (Table S1). We restricted the phylogenetic placement of the 194 taxa without morphology by using genus-level monophyletic clade constraints. Since all genera in our tree are represented in our morphological matrix, all species could be placed within a monophyletic generic lineage with at least one member of that genus having morphological information.

To make direct and computationally tractable comparisons, we focused the analyses of posterior tree distributions on the output from the resolved and semi-resolved analyses with the exponential origin prior. We first pruned the 194 species without morphology from the semi-resolved analysis trees, so that trees from both analyses had the same leafset, then randomly subsampled the distributions to 900 trees each (1% of the post-burnin tree sample). We calculated stratigraphic congruence metrics for each tree in each posterior distribution using the R package ‘strap’ (Bell and Lloyd, 2015). We used the stratigraphic consistency index (SCI; (Huelsenbeck, 1994), minimum implied gap (MIG; (Norell and Novacek, 1992)), and gap excess ratio (GER; (Wills, 1999).

To visualize the effects on topology and, consequently, stratigraphic congruence, we also produced a treespace of these trees from both posterior distributions using the R package ‘TreeDist’ (Smith, 2020b). Further, we used the treespace to produce a stratigraphic congruence ‘landscape’. We also explored leaf stability through the effect of ‘rogue’ taxa using the ‘Rogue’ R package (Smith, 2022a). Finally we assessed the quality of consensus trees by calculating the amount of information (bits) they communicated about clades in the posterior distribution using the splitwise phylogenetic information content (SPIC; (Smith, 2022a).

A detailed explanation of the dataset, analytical setups, convergence assessment, and analysis of posterior distributions and stratigraphic congruence are provided in the Supplementary Methods. R code for writing Beast XML files with large amounts of PDBD information and taxonomic constraints, and all analyses of the posterior tree distribution, are available in the Supplementary Material.

## Results

### Stratigraphic congruence and topology

The semi-resolved FBD analysis was superior in all measures. Semi-resolved analyses produced trees that were significantly and substantially more stratigraphically congruent under all metrics (Fig. 1): a consequence of different topologies in the posterior tree distributions. The effect is more pronounced in the MIG and GER than the SCI, which is a consequence of the SCI ignoring implied gaps (i.e., ghost ranges). When analysed together, trees from each analysis largely occupy separate regions of treespace (at least in the first dimension). Consequently, they occupy distinct regions of the stratigraphic congruence tree ‘landscape’, with a clear gradient of congruence (Fig. 2A; Fig. S2).

**Figure 1.**
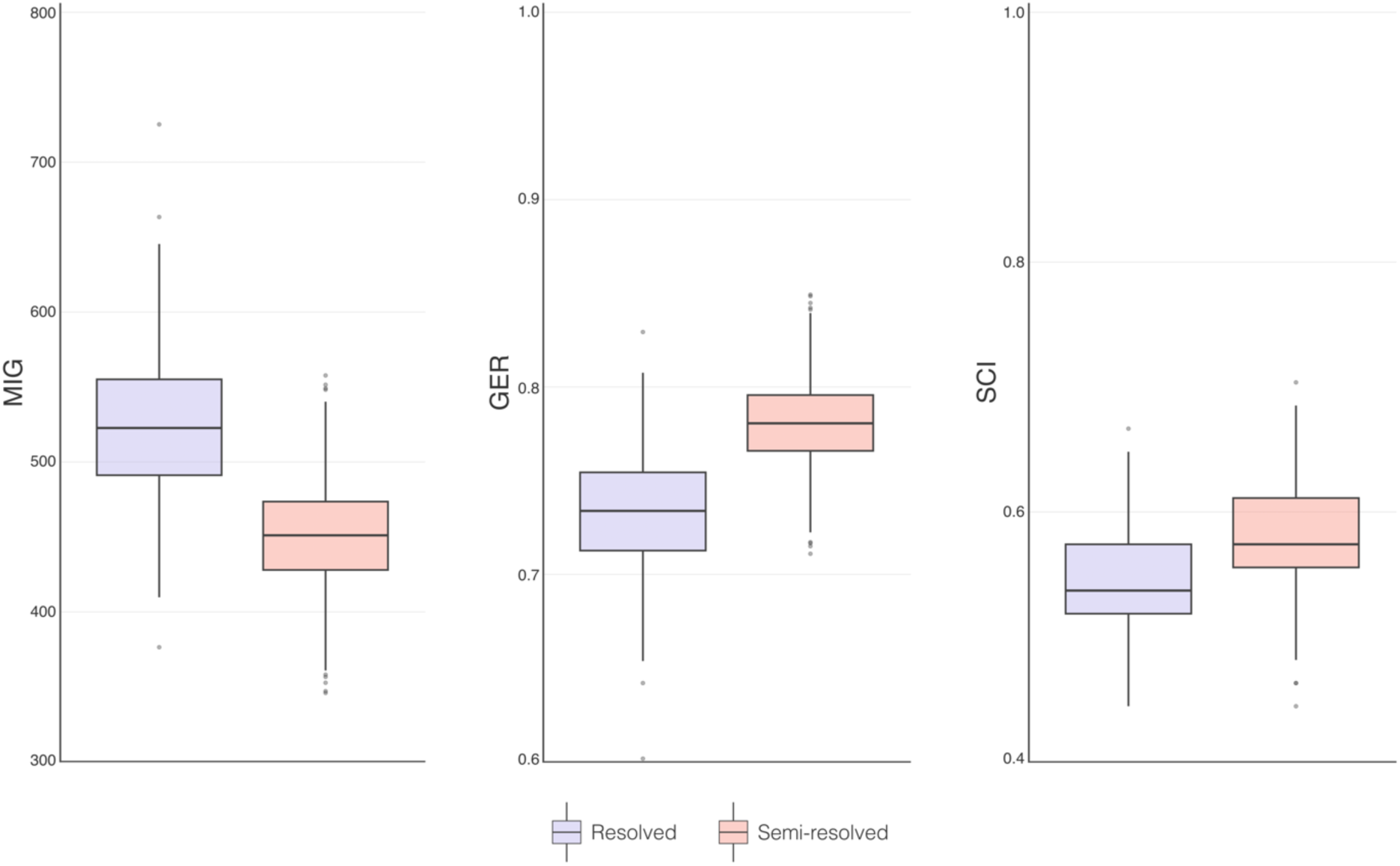
Stratigraphic congruence metrics for the posterior distributions of trees produced from resolved and semi-resolved FBD analyses. MIG = minimum implied gap (lower is better), GER = gap excess ratio (higher is better), SCI = stratigraphic consistency index (higher is better). All pairwise p values <0.05.

**Figure 2.**
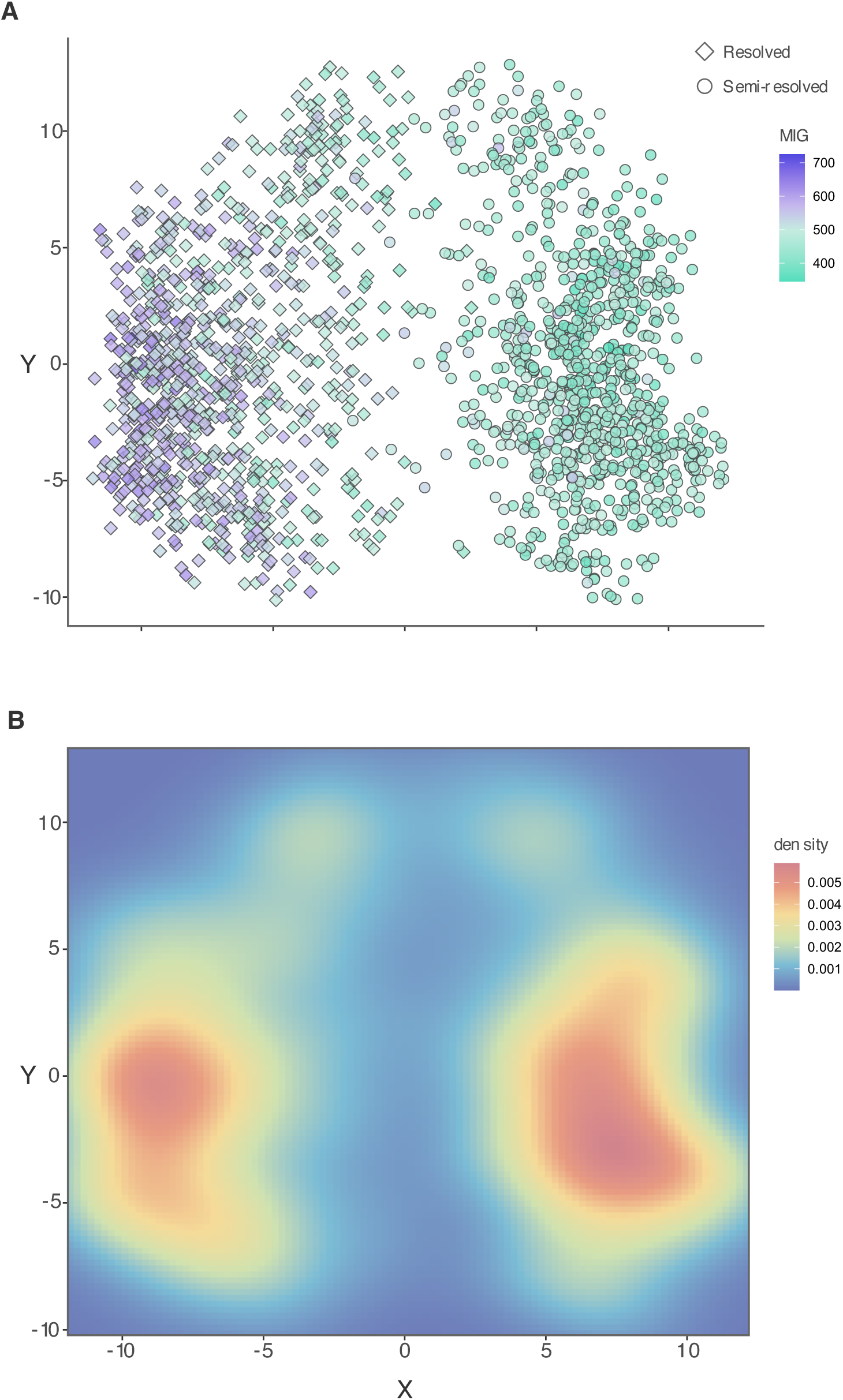
Treespace produced from subsampled trees from both analysis types. A) Stratigraphic congruence ‘landscape’ (points coloured by MIG; lower values indicate higher stratigraphic congruence). Analysis type is represented by point shape. B) Heatmap of density in the same treespace. Warmer colours indicate a higher density of trees clustering in that region of treespace.

There were important differences in the spread of each distribution in treespace. Trees from the semi-resolved analyses clustered more densely around the region of highest stratigraphic congruence in the landscape (Fig. 2B). Meanwhile, trees from the resolved analyses appeared largely on a local optimum of lower stratigraphic congruence and were also more diffuse. This is reinforced by the semi-resolved distribution having lower sum of variances, sum of ranges and mean centroid distance (Table S3). While topological differences are apparent in the first dimension of the treespace, the distributions overlap in other dimensions (Fig. S3).

### Consensus/summary trees and leaf stability

Node supports were relatively low for some parts of the tree in both analyses, but key parts of the tree had higher support in the semi-resolved analysis. Additionally, the sum of posterior probabilities for all nodes in the maximum clade credibility (MCC) tree from the semi-resolved analysis was higher than for the resolved analysis (Table 1). Semi-resolved analyses also produced slightly more resolved majority rule consensus trees, having one more node than the resolved analysis consensus tree (Table 1; Fig S4).

**Table 1.** Statistics related to consensus trees (majority rule and maximum clade credibility) for the posterior distributions produced from the resolved and semi-resolved FBD analyses. Statistics concern the majority rule consensus tree, except for the sum of posterior probabilities, which concerns the MCC tree.

| Analysis | N nodes | N rogues | Information content | Mean improvement from rogue removal | Sum of posterior probabilities (MCC tree) |
| --- | --- | --- | --- | --- | --- |
| Resolved | 20 | 12 | 129.38 | 6.58 | 18.96 |
| Semi-resolved | 21 | 6 | 185.59 | 5.49 | 21.68 |

The semi-resolved analysis consensus tree also had a higher information content (SPIC) than did the resolved analysis consensus tree (Table 1). Higher information content means the semi-resolved analysis consensus captured deeper nodes/larger clade relationships, reflecting greater stability of those relationships in the posterior tree distribution.

Rogue taxa are those taxa whose phylogenetic position is particularly ‘unstable’ in a distribution of trees. These taxa create issues in summarising tree distributions. Here, the resolved analysis had a posterior distribution with double the taxa displaying rogue behaviour than the semi-resolved analysis. Further, each rogue in the resolved distribution was more detrimental to the consensus tree on average than were the semi-resolved consensus rogues. The removal of rogue taxa also had the effect of increasing the MIG in both analyses (Fig. S5).

### Divergence and origin times

Estimates of the origin age were more precise with the semi-resolved analysis than with the resolved analysis (Fig. 3A). The 95% highest posterior density (HPD) interval for the origin age in the semi-resolved analysis was 25.7% narrower with a uniform prior and 31.3% narrower with an exponential prior than the resolved analysis. The HPDs were almost entirely overlapping, demonstrating the semi-resolved analysis mainly produced a marked decrease in the uncertainty. Notably, however, the median inferred ages were slightly younger for the semi-resolved analyses.

**Figure 3.**
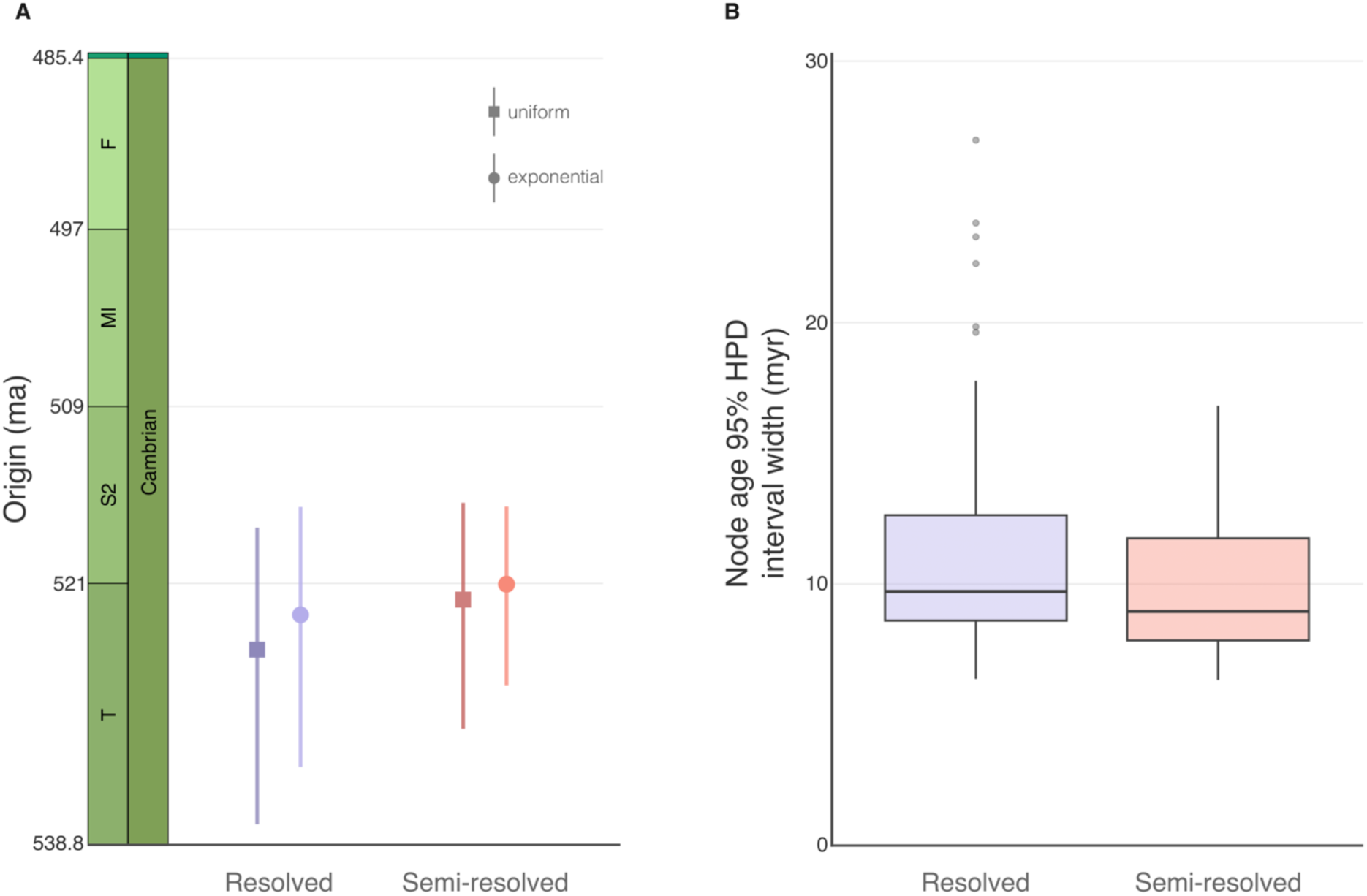
Comparison of effects of resolved and semi-resolved FBD analyses on inferred origin and divergence times. A) 95% highest posterior densities (HPDs) and medians of the origin time for each type of analysis and prior distribution (square = uniform; circle = exponential). B) 95% HPD width for all nodes in each MCC tree.

Inferred clade divergence times were also markedly more precise for the semi-resolved analysis than the resolved analysis (Fig. 3B). The largest uncertainty interval around a node for the resolved analysis was 27 myr, while it was 17 myr for the semi-resolved analysis.

## Discussion

Our comparative analyses show that a combined analysis including both fossils with and without character data using taxonomic constraints is not only possible in practice but outperforms analyses of only taxa with morphology for fossil groups. As expected, more fossil data and a more complete representation of their temporal distribution (as in the ‘semi-resolved’ analysis) produces markedly more precise parameter estimates. Any analyses where divergence time estimation is of interest would therefore benefit from the combined ‘semi-resolved’ approach. Also unsurprisingly, trees produced from the ‘semi-resolved’ analysis show much higher levels of stratigraphic congruence than those from the ‘resolved’ analysis. That is, they better fit the order of appearance of taxa in the fossil record. This may be an intuitive result, but it illustrates how sampling only taxa with abundant morphological characters has the potential to generate misleading results.

Stratigraphic age information for taxa with morphology has previously been shown to improve phylogenetic analyses of those taxa (Barido-Sottani et al., 2023; Mongiardino Koch et al., 2021). In such analyses, age information causes some regions of treespace to become implausible. Moreover, topological changes to regions of the tree with weaker character signal can be induced by age information (King, 2021). A crude way to visualise these changes is by simply comparing the MCC trees and the ghost ranges (Fig. S6). However, MCC trees are often unreliable for tree distributions with high levels of uncertainty, as is the case for most morphological datasets (O’Reilly and Donoghue, 2018). Understanding and comparing the posterior distributions is therefore aided here by the visualisation of treespace, a powerful tool for exploring distributions of trees (Smith, 2022b), but underutilised particularly for fossil groups (Wright and Lloyd, 2020). In this treespace, trees from the ‘semi-resolved’ analysis did not occupy the more stratigraphically incongruent regions, and clustered more densely around the regions of highest stratigraphic congruence. Our results therefore show that age information of taxa other than just those with morphology can further cause stratigraphically incongruent regions of treespace to become implausible.

The ‘semi-resolved’ analysis had a significant and substantial effect on the SCI, but had a larger effect on the MIG and GER metrics. The SCI is a proportion of stratigraphically consistent nodes in a tree, and does not consider the length of ghost ranges, like the MIG and GER do (Bell and Lloyd, 2015). This suggests that the restructuring of only a few nodes (or taxa) to stratigraphically consistent positions can break up long branches. This is a direct consequence of a more representative distribution of fossil sampling ages.

Leaf stability is another important aspect of a distribution of topologies. Taxa whose phylogenetic position are particularly unstable throughout a distribution of trees – ‘rogues’ – are problematic as they blur otherwise stable phylogenetic relationships (Nixon and Wheeler, 1992; Pohle et al., 2022; Smith, 2022a). They also obliterate node support values and the ability to resolve relationships in consensus trees. Here, the ‘semi-resolved’ produced fewer rogue taxa. It may be that the additional constraints provided by the stratigraphic information prevent would-be rogues from attaching to very distant, stratigraphically inconsistent, branches. This would also align with the aforementioned observation that age information exerts strongest effects in sections of the tree with weaker character signal (King, 2021). Nevertheless, this higher leaf stability affords more confidence in the inferred clade relationships than the ‘resolved’ analysis.

Perhaps due to fewer rogue taxa obscuring the resolution of deeper nodes, the semi-resolved analysis produced a majority rule consensus tree with higher information content than the resolved analysis. Splits in a summary tree are not all equally informative, for example, a split leading to two species in the same genus is less informative than a split separating two families. While intuitive, formally, this is because the probability of larger clades and splits among them is much lower in a posterior distribution. When a consensus tree captures relationships of lower probability in the tree distribution it is therefore more informative (Shannon, 1948; Smith, 2022a; Steel and Penny, 2005). Consequently, majority rule consensus trees with the same number of resolved splits may not necessarily be equally informative. Here, the higher information content in the semi-resolved majority rule consensus tree is a consequence of it capturing larger clade relationships at deeper nodes (Fig. S4), and thus better summarising the posterior distribution. Put another way, this reflects a higher level of clade stability in the posterior distribution.

The combined approach to incorporating fossils into FBD analyses, here termed ‘semi-resolved’, offers an intuitive and relatively simple means of mitigating taxon sampling biases, while providing a more comprehensive utilisation of the fossil record. Yet, this strategy has historically been overlooked. We tested and demonstrated the effects of this approach by simply incorporating bulk occurrence stratigraphic age information from the PBDB into FBD phylogenetic analyses of a fossil group. This information is readily available and easily obtained, so may offer a high benefit:effort ratio. Nevertheless, occurrences must be assigned to the tree with caution. (Barido-Sottani et al., 2023) show that erroneously placed fossil taxa can not only nullify the positive effects but reduce overall accuracy in phylogenetic inference. Appropriately assigning taxonomic constraints to a dataset/analysis therefore requires a taxonomic expert. Moreover, while the results we present are encouraging, we echo sentiments in (Barido-Sottani et al., 2023) that developments in phylogenetic and paleobiological models are no replacement for fundamental taxonomic, stratigraphic and systematic research.

## Conclusion

We corroborate previous simulation results (Barido-Sottani et al., 2023) with a complex empirical scenario, and further emphasise the utility of stratigraphic age information in phylogenetic analyses. While the focus here was on a group of extinct organisms, our findings would also be applicable to groups including extant taxa. We therefore encourage researchers to consider the full amount of fossil information that fundamental taxonomic and stratigraphic work has afforded.

## Supplementary material

### Supplementary methods

#### Overview of analytical setups

To address each of our questions and test sensitivity to certain priors, we set up four different analyses (Table S1). The differences in each analysis are the inclusion/exclusion of taxa with age information only (i.e., no morphological information) (resolved vs semi-resolved FBD) and type of prior on the origin age.

#### Taxa and Character matrix

We used the character matrix for trilobites from a forthcoming publication detailing phylogenetic analyses and subsequent revision of a trilobite order. The matrix is deposited on Morphobank and can be found there (http://morphobank.org/permalink/?P5740). Taxa in this matrix were sampled to address questions of higher taxonomic classification in trilobites and an associated so-called “cryptogenesis” problem. This problem refers to a lack of understanding of the phylogenetic relationships between Cambrian and post-Cambrian groups (Paterson, 2020). Selection of included taxa was also justified on the basis of, where possible, having well described or represented material, including as much of the morphology as possible, as well as those taxa with known articulated specimens. These taxa span ∼125 ma, from the Middle Cambrian to the Middle Devonian.

The matrix includes 56 taxa and 254 discrete characters, resulting in the largest trilobite character matrix yet (with regard to characters), and employs a hierarchical/contingent coding strategy to best accommodate inapplicable characters. Characters are binary or multistate and continuous characters were discretised.

#### Temporal data

Temporal information associated with each taxon in the morphological matrix was based on the finest stratigraphic interval in which the fossil material used for the original description was recovered (Supplementary Material). For the majority of taxa, this was a regional (trilobite, agnostid, conodont, or graptolite) biozone, which we correlated to a global scale based on (Gradstein et al., 2020) to obtain absolute ages. For a few taxa, only the formation could be used for correlation and in rare cases only a position within a regional stage was available, resulting in larger amounts of uncertainty when correlating to the global scale.

While we had temporal information for every one of the taxa included in the morphological matrix, our major aim was to assess the effects of a considerable increase in amount of fossil information in an empirical setting. In service of this, we used the Paleobiology Database (PBDB; https://paleobiodb.org) to retrieve records of occurrences from the families of the taxa included in our morphological matrix (Supplementary Material). We examined and verified the quality of dataset retrieved from the PBDB to determine the most appropriate taxonomic level to use in our analyses.

The raw downloaded dataset contained many aberrant age ranges for taxon occurrences (e.g., ∼100 million years; Fig. S1A) and spurious and/or uncertain taxonomic assignments. To avoid these issues, we filtered the dataset to only include those occurrences that were identified to species and were within the genera included in the morphological matrix. This generated a dataset with much more refined and acceptable age ranges, and mitigated the effects of erroneous taxonomic placement when using topological constraints (Barido-Sottani et al., 2023a). This dataset included 793 occurrences and the median number of occurrences for each species was one. Many taxa, however, had multiple occurrences in the same time interval, which did not affect the upper and lower age bounds for that taxon. Of the 275 unique occurrences, some taxa had occurrences with different age intervals/ranges. For these 53 taxa with multiple unique occurrences, we therefore randomly, uniformly subsampled a single occurrence for each taxon, such that each taxon and its age interval was represented by a single fossil. We used the random subsampling approach to meet the FBD model assumption of uniform/random sampling as closely as possible while ensuring our analyses remained computationally feasible. Selecting intervals based on the first appearance, for example, results in biased and inaccurate results (Matschiner et al., 2017).

Some occurrences were still associated with intervals that were relatively poorly resolved, reflecting large amounts of uncertainty (e.g., crossing multiple stage boundaries). Still, we considered these to be acceptable, as age estimates of fossils from poorly dated/resolved deposits are still relatively accurate when inferred with the FBD model (Barido-Sottani et al., 2023b). This is especially so when such unresolved ranges are few and the proportion of highly resolved ages is high, as is the case in our dataset.

Finally, we removed one species from this dataset that had a range interval as “Cambrian” and was based on an outdated geologic time scale which placed its upper bound in the Ediacaran. Occurrences of taxa in the morphological matrix were also removed, as these had already been obtained from the literature. This resulted in a dataset of 194 taxa with age uncertainty intervals (Fig S1B), and a combined dataset of 250 taxa.

#### Phylogenetic inference

For all of our analyses, we use the fossilized birth-death model (FBD), as it offers the most appropriate means to incorporate vast amounts of fossil-specific information. We conducted all analyses in the software Beast 2.6.7 (Bouckaert et al., 2019), using the constant rates fossilized birth-death (FBD) model as implemented in the Sampled Ancestors package (Gavryushkina et al., 2014). As opposed to diversification, turnover rates, and sampling proportion as is commonly used as the default in Bayesian phylogenetic software, we parametrized the FBD model using the “canonical” parameterisation with speciation (birth; α), extinction (death; μ) and sampling (ψ) rates. The only changes among analyses were the included taxa (and associated age information) and the prior on the origin time, otherwise all parameters and priors remained constant.

##### FBD model parameters

We put a lognormal distribution on both birth and death rates with means of 0.022 and 0.029 respectively, based on estimates from empirical data, and both with a standard deviation of one. We estimated appropriate starting values and prior distributions for speciation and extinction rates of all trilobites through the Paleozoic from the PBDB. To do this, we sampled-standardized the data with shareholder quorum subsampling (Alroy, 2010) and calculated mean per capita rates per million years. The sampling rate prior was lognormal distribution with a mean of 0.3 and standard deviation of 0.2. We converted the per-interval sampling probability for trilobites given in (Foote and Sepkoski Jr, 1999) to the rates used here.

##### Morphological model

For the model of morphological character evolution, we used the Mkv model with the correction for variable character ascertainment bias (Lewis, 2001). We partitioned the characters by number of states, so that each character had an appropriate Q-matrix, and allowed for rate heterogeneity among characters with four discrete gamma rate categories (Yang, 1994). We also placed an exponential prior with a mean of one on the alpha parameter of the gamma distribution to estimate it. To reduce computational complexity and unnecessary overparameterization, we shared the gamma rate categories among all partitions.

##### Clock model

As a clock model, we used a relaxed lognormal clock, with an unbounded uniform distribution on the mean rate, and a gamma distribution on the standard deviation with the alpha parameter set at 0.540 and the beta parameter set at 0.382.

##### Origin prior

While we did not attempt to investigate the origins of the entire trilobite group – and the taxa we did investigate are not its earliest members – we assessed the effects of the increased fossil information between two origin time prior types. In doing so, we also tested the sensitivity of origin age estimation to two different priors (summarised in Table S1).

Based on a tip-dated phylogenetic analysis of Cambrian trilobites, (Paterson et al., 2019) suggest a conservative origin age for all trilobites around the Ediacaran-Cambrian boundary, around 541 ma, with a median estimate of ∼535 ma. Note that this was based on a now outdated estimate of the age of this boundary, which has since been altered to 538.6 - 538.8 ma as inferred in (Linnemann et al., 2019). Contrarily, (Holmes and Budd, 2022) suggest the fossil record is a more faithful representation of the evolutionary history of trilobites, and as such, the true origin age should lie somewhere very close to the first appearance of trilobite body fossils around 521 ma.

We used a uniform prior with an upper bound around the Ediacaran-Cambrian boundary (538.5 ma) and a lower bound at the lower bound of the uncertainty interval of the oldest taxon in the analyses. Considering a possible origin much closer to the first appearance of fossils in our analyses, we also use an exponential prior, with the same bounds as the uniform prior. This places much higher probabilities on origin ages close to the age of the first sampled tip, which decay exponentially with time.

##### Taxonomic constraints and ages

Because the 194 fossil species were not associated with morphological information here, we used genus-level monophyletic clade constraints to restrict their phylogenetic placement. Therefore, all species would be placed within a monophyletic generic lineage with at least one member of that genus having morphological information, since all genera in our tree are represented in our morphological matrix.

We fixed the age of the youngest taxon at 0 and scaled ages of all other taxa accordingly. In the 250-taxon dataset, multiple taxa were equally youngest, therefore we sampled one taxon at random to be fixed to 0. Post-scaling, taxa were then assigned a uniform distribution between the bounds of their associated temporal interval to sample their age.

##### Analysis

For each analysis, we ran two independent chains of 1,000,000,000 or 2,000,000,000 MCMC generations, sampling every 10,000 or 20,000, for the ‘resolved’ and ‘semi-resolved’ analyses respectively, discarding the first 10% as burn-in. We assessed parameter convergence in Tracer (Rambaut et al., 2018) and also determined whether the independent chains had converged on the same posterior distribution. The parameter log and tree files from independent runs were combined and resampled every 20,000 or 40,000 states. Then finally, we generated maximum clade credibility (MCC) trees created in TreeAnnotator (Bouckaert et al., 2019).

#### Analysis of posterior distributions

For the following analyses, we focused on the output from the posterior distributions produced from the resolved and semi-resolved analyses with exponential priors on the origin. We did this to keep comparisons direct and computationally tractable, and because the differences in posteriori distributions between the uniform and exponential prior analyses were relatively minor.

The taxa without morphology were only involved in informing parameter (including topology) estimation of the FBD analysis and were no longer necessary in the trees. As such, we first pruned all taxa without morphological information from the posterior distribution of trees. This also ensured that both distributions had trees with the same leafset and direct comparisons could be made between them. For computational tractability, we then resampled the distributions to 900 trees (1% of the 90,000 post-burnin sampled tree distribution).

##### Stratigraphic congruence

We calculated stratigraphic congruence metrics for each tree in the posterior distributions using the R package ‘strap’ (Bell and Lloyd, 2015). We used the stratigraphic consistency index (SCI; (Huelsenbeck, 1994), minimum implied gap (MIG; (Norell and Novacek, 1992)), and gap excess ratio (GER; (Wills, 1999). Briefly, SCI measures the proportion of nodes in a tree for which the descendants of that node are the same age or younger than the oldest sister taxon of that node, i.e., it is stratigraphically consistent. As a proportion, its values can range from 0 to 1, where 1 represents a tree whose nodes are perfectly ordered stratigraphically. Note, however, that it is purely a proportion of nodes and therefore does not consider the size of implied gaps (i.e., ghost ranges). MIG is a measure of the total ghost ranges in a tree. GER is an extension of the MIG, which subtracts the best possible stratigraphic fit from the MIG, then scales this by the difference between best and worst possible fit values. See Bell and Lloyd (2015) for a more detailed review of each of these metrics.

##### Treespace

To visualize the distributions of trees produced from the different analyses and highlight any differences resulting from the inclusion of the additional fossil information, we generated a treespace of the tree distributions from both analyses. Treespaces are produced through mapping metric distances among trees in high-dimensional spaces, then using ordination, or multidimensional scaling (MDS), techniques to represent these spaces in fewer dimensions. Several distance metrics exist, but here we use the clustering information (CI) distance (Smith, 2020b) implemented in the R package ‘TreeDist’ (Smith, 2020a, p. 2). The CI distance is a type of generalized Robinson-Foulds (RF) distance (Robinson and Foulds, 1981) that also accounts for the differing information content in different sized splits. Moreover, it recognizes where pairs of splits are similar, but not exactly equivalent; this gives it the added advantage of also being much less negatively affected by rogue taxa, while also being relatively fast to calculate. Given these desirable properties, the CI distance provided a suitable distance metric and classical MDS to produce our treespace. Finally, we produced a tree ‘landscape’ (St. John, 2017) by colouring each point (tree) in the treespace by its stratigraphic congruence.

##### Consensus and rogue taxa

Interrogating the majority rule consensus trees allowed us to compare the major topological differences and further assess the impact of the different analysis types on inferred topologies and leaf and clade stability. Here, stability refers to how frequently a clade is sampled in the posterior distribution. Stable leaves produce stable clades when their position relative to related members does not change much in the posterior sample.

We first assessed the resolution of consensus trees by simply calculating the number of nodes. A greater number of nodes in a majority rule consensus tree should reflect greater stability of clades in a posterior distribution of trees. That is, that those clades would be stable and robust enough to occur in 50% or greater of the trees in the posterior distribution. However, not all nodes are equally informative: the node resolving the relationship between two members of the same genus is not as informative as the node resolving the relationship between two families, for example. To quantify the informativeness of relationships resolved in the consensus trees, we calculated the splitwise information content (SPIC) of each consensus tree using the ‘Rogue’ R package (Smith, 2022a)

We also explored the effects of leaf stability by identifying taxa displaying rogue behaviour in both posterior tree distributions using the ‘Rogue’ R package (Smith, 2022a). We calculated the mean improvement of successively removing rogue taxa to determine the average impact each rogue taxon had on the consensus tree. Finally, to assess the influence of rogue taxa on stratigraphic congruence, we pruned the rogue taxa that had an outsized effect on consensus information content from the posterior distribution and recalculated stratigraphic congruence metrics for these trees.

R code for the entire analytical pipeline can be found in the Supplementary Material. This includes code to extract prior parameter estimates, subsample fossils from the larger PBDB dataset, a function and script to generate Beast XML files with large numbers of taxonomic constraints and age priors, and all code to analyse the posterior distributions.

---

**Table S1.**
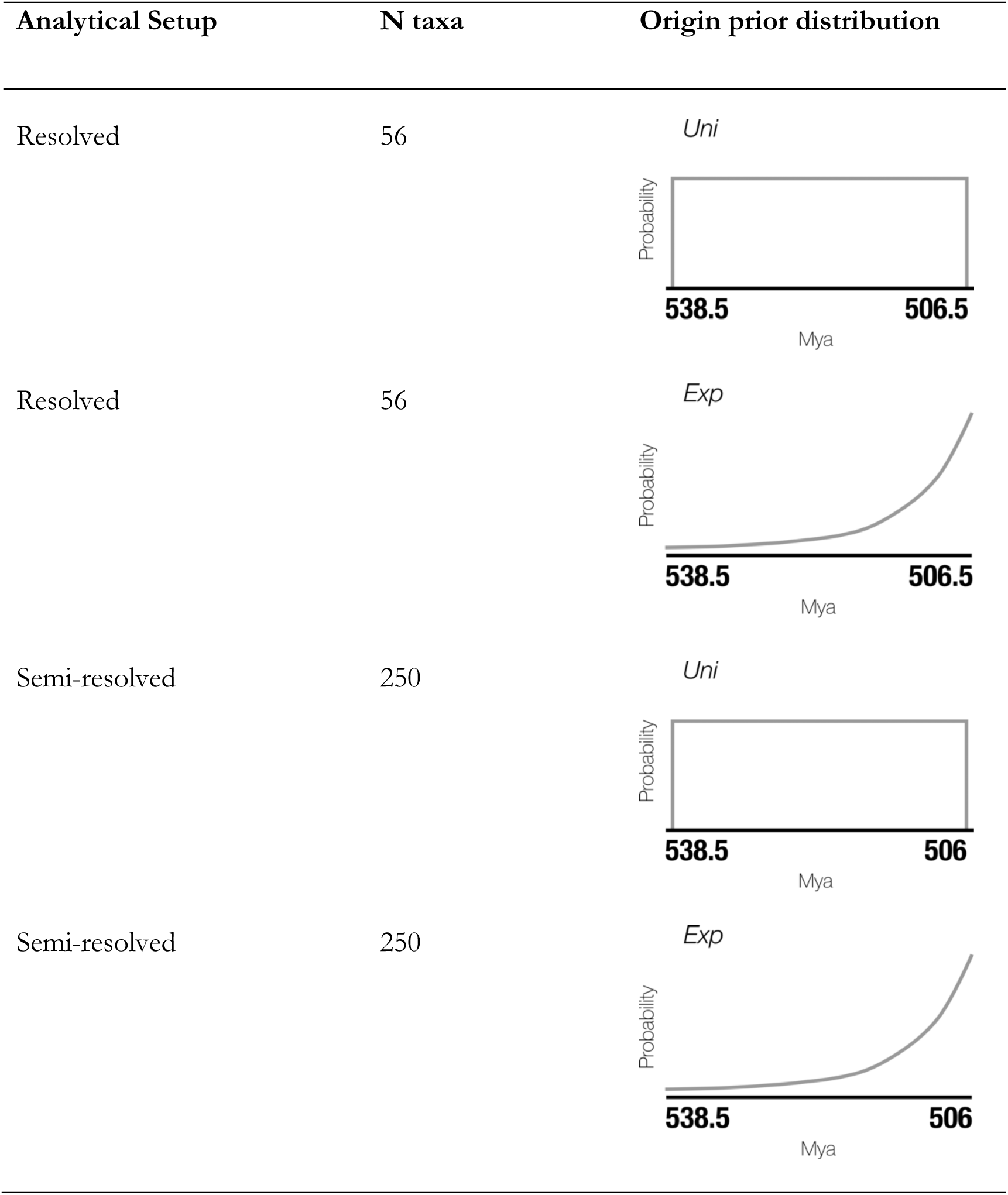
Details of analytical setups used in this study.

**Figure S1.**
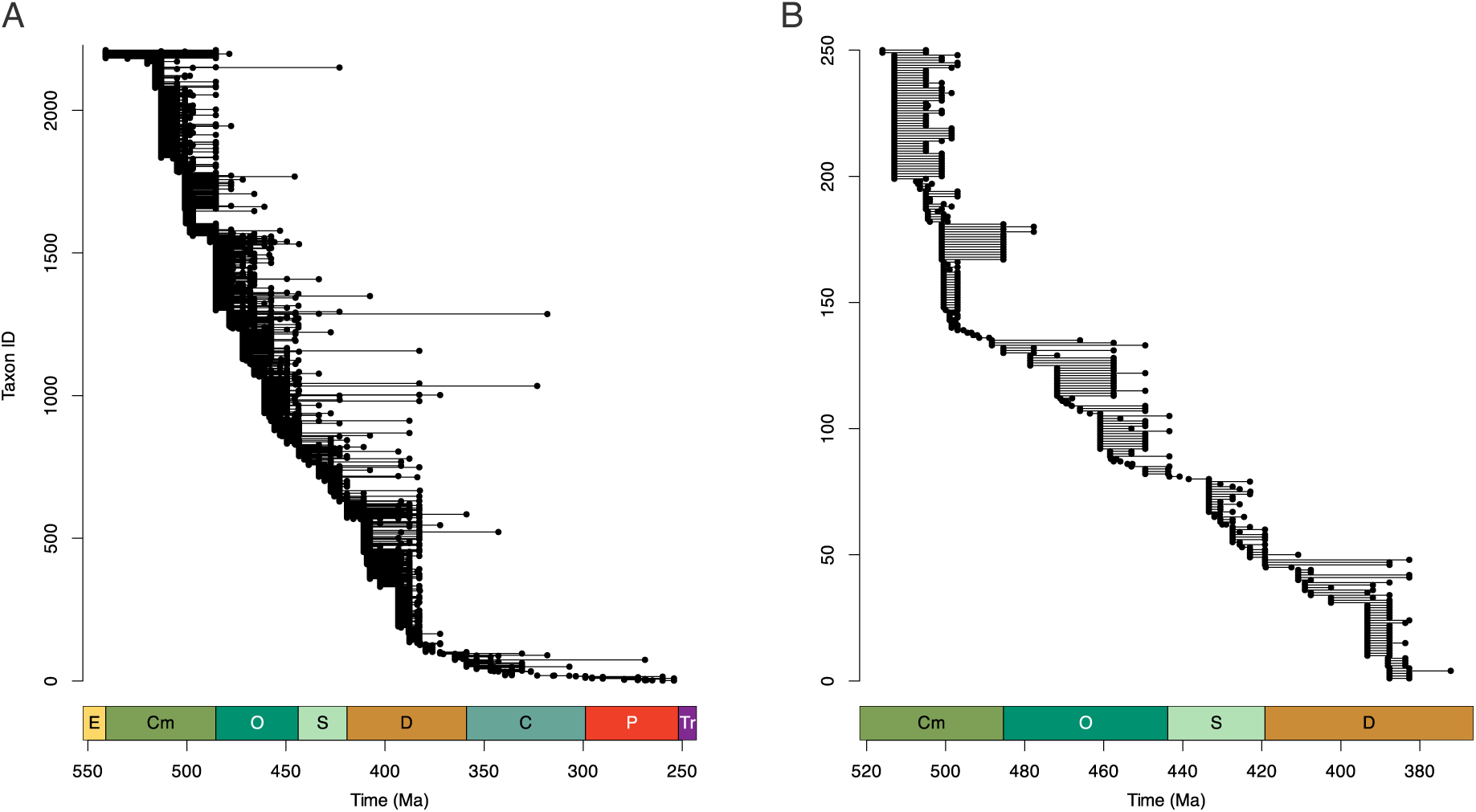
Taxon ranges from from the raw PBDB dataset (A) and the cleaned dataset used for the analyses (B).

## Supplementary table and figures

**Table S2.** Measure of spread/dispersion of each posterior distribution in treespace.

| Analysis | Sum of variances | Mean centroid distance | Sum of ranges |
| --- | --- | --- | --- |
| Resolved | 34.59 | 5.21 | 37.24 |
| Semi-resolved | 30.48 | 4.78 | 37.02 |

**Figure S2.**
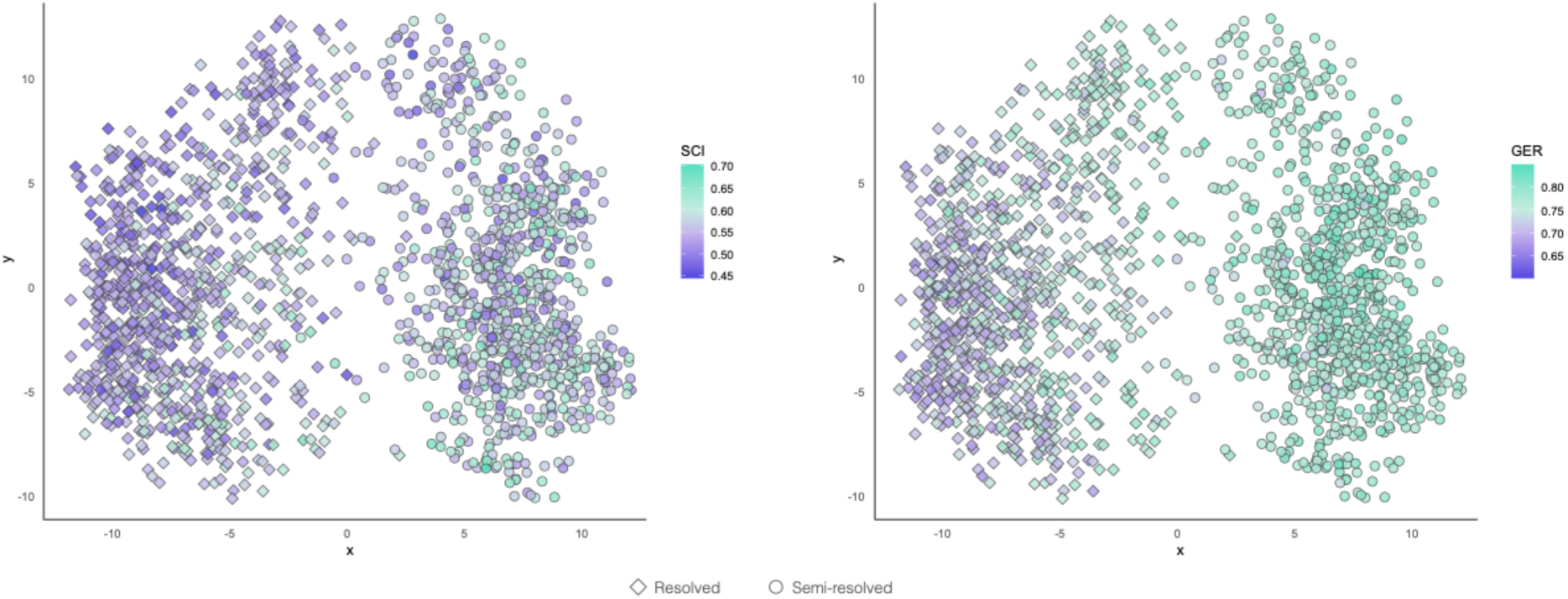
Tree ‘landscapes’ of stratigraphic congruence by SCI and GER (higher is better for both metrics). Analysis type is represented by point shape.

**Figure S3.**
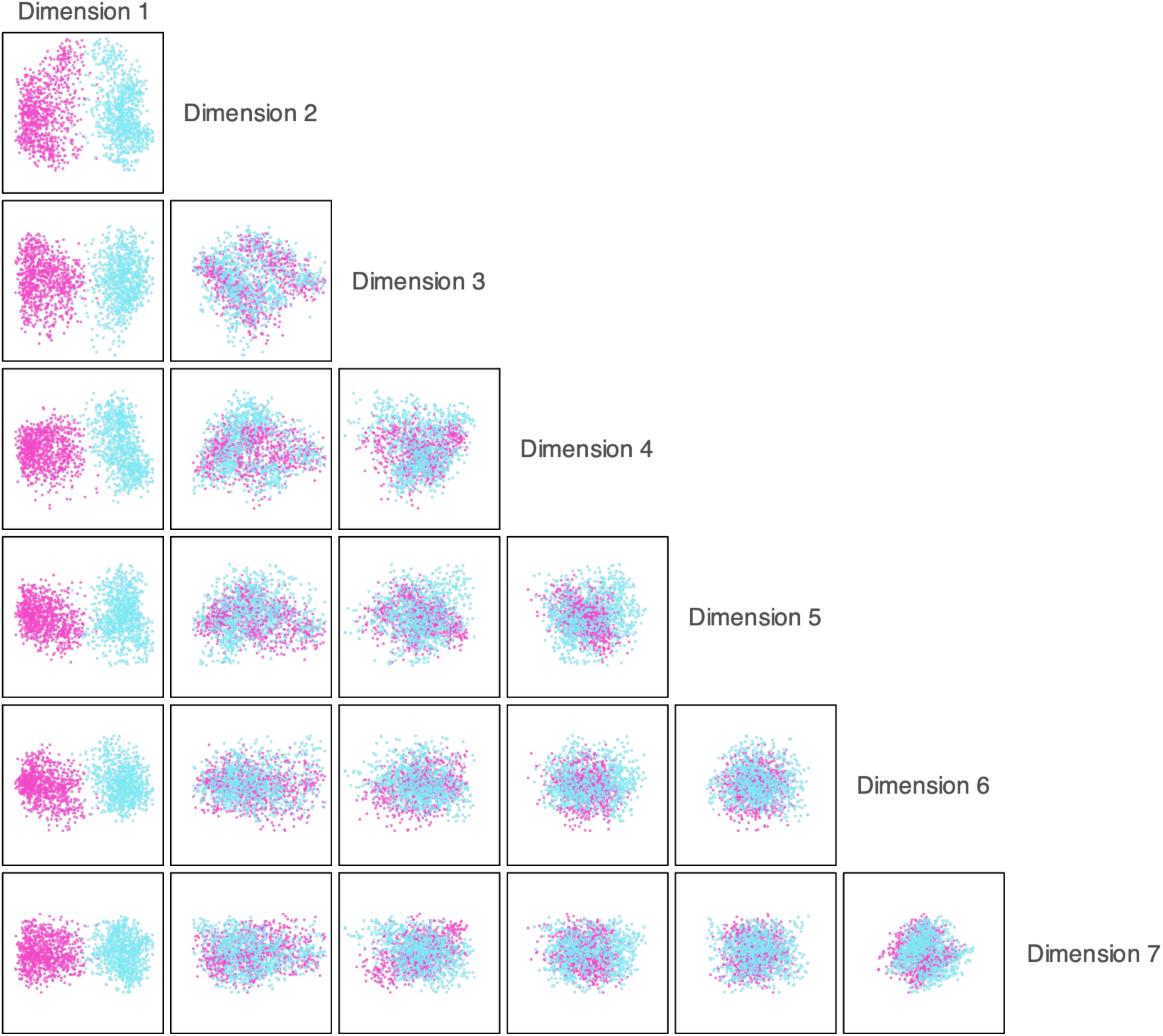
Higher dimensions of the treespace of the posterior tree distributions of both analysis types. Pink = resolved, blue = semi-resolved.

**Figure S4.**
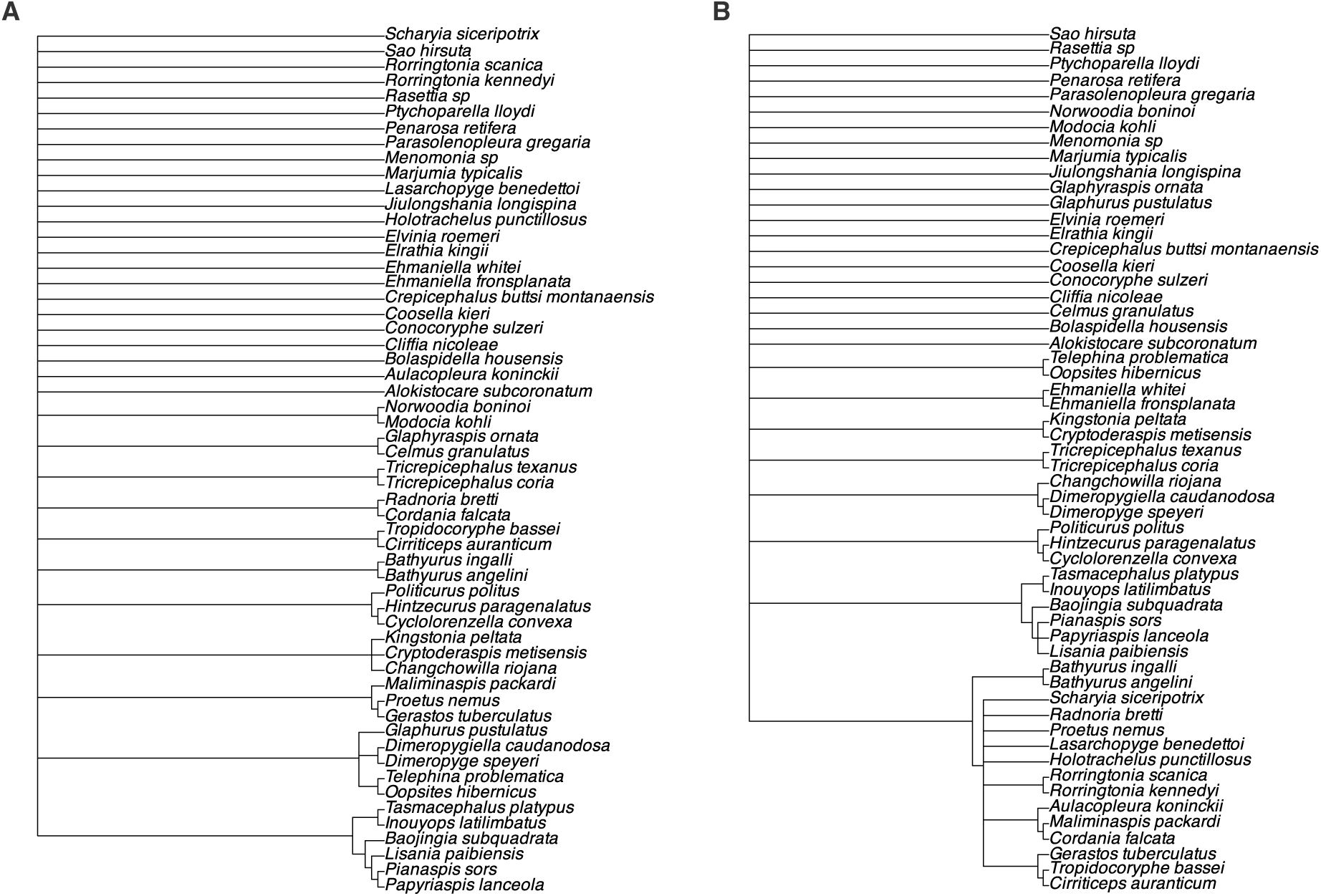
Majority rule consensus trees from the resolved (A) and semi-resolved (B) analyses.

**Figure S5.**
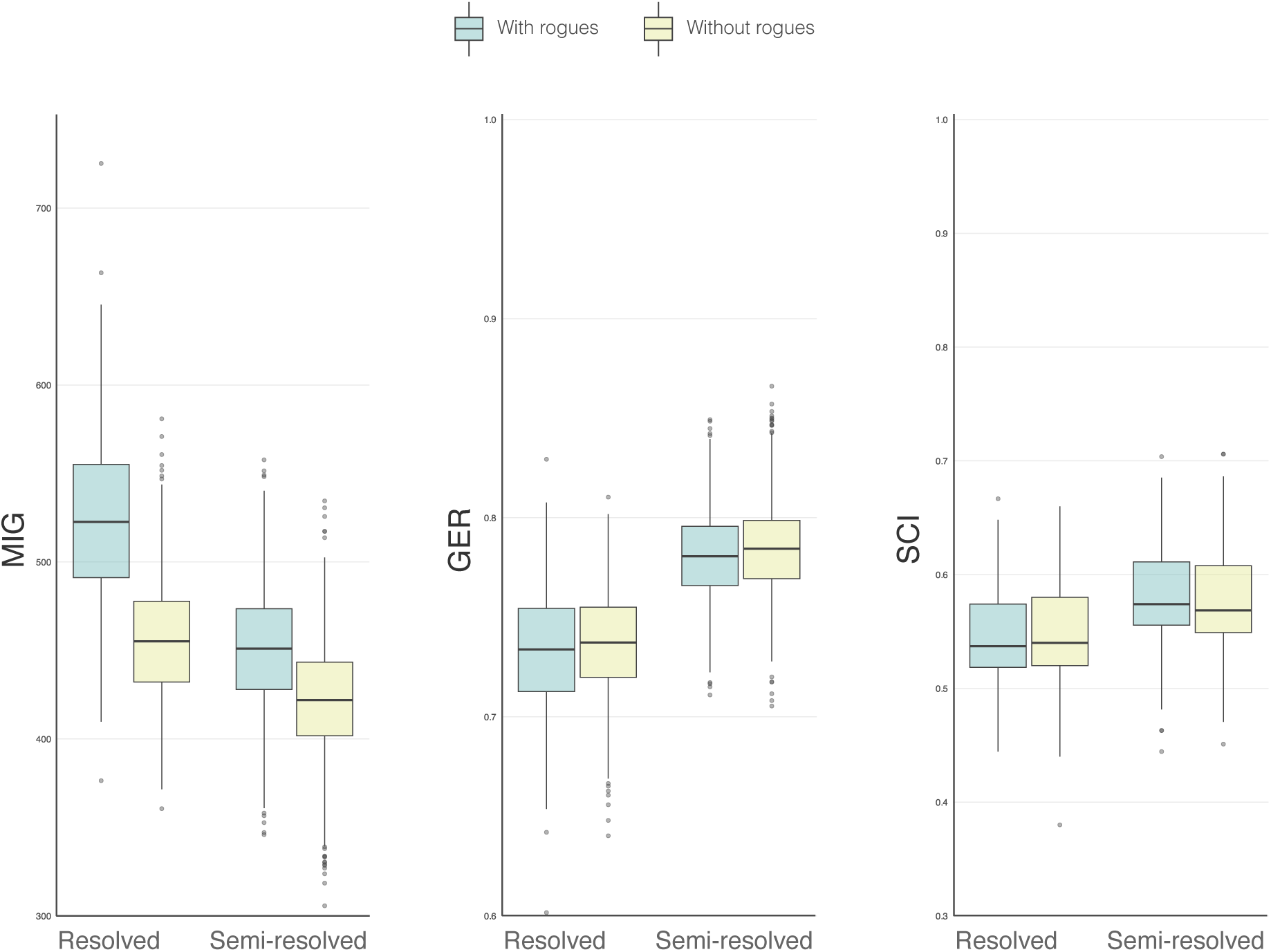
Stratigraphic congruence of the posterior distributions from both analysis types with and without rogues.

**Figure S6.**
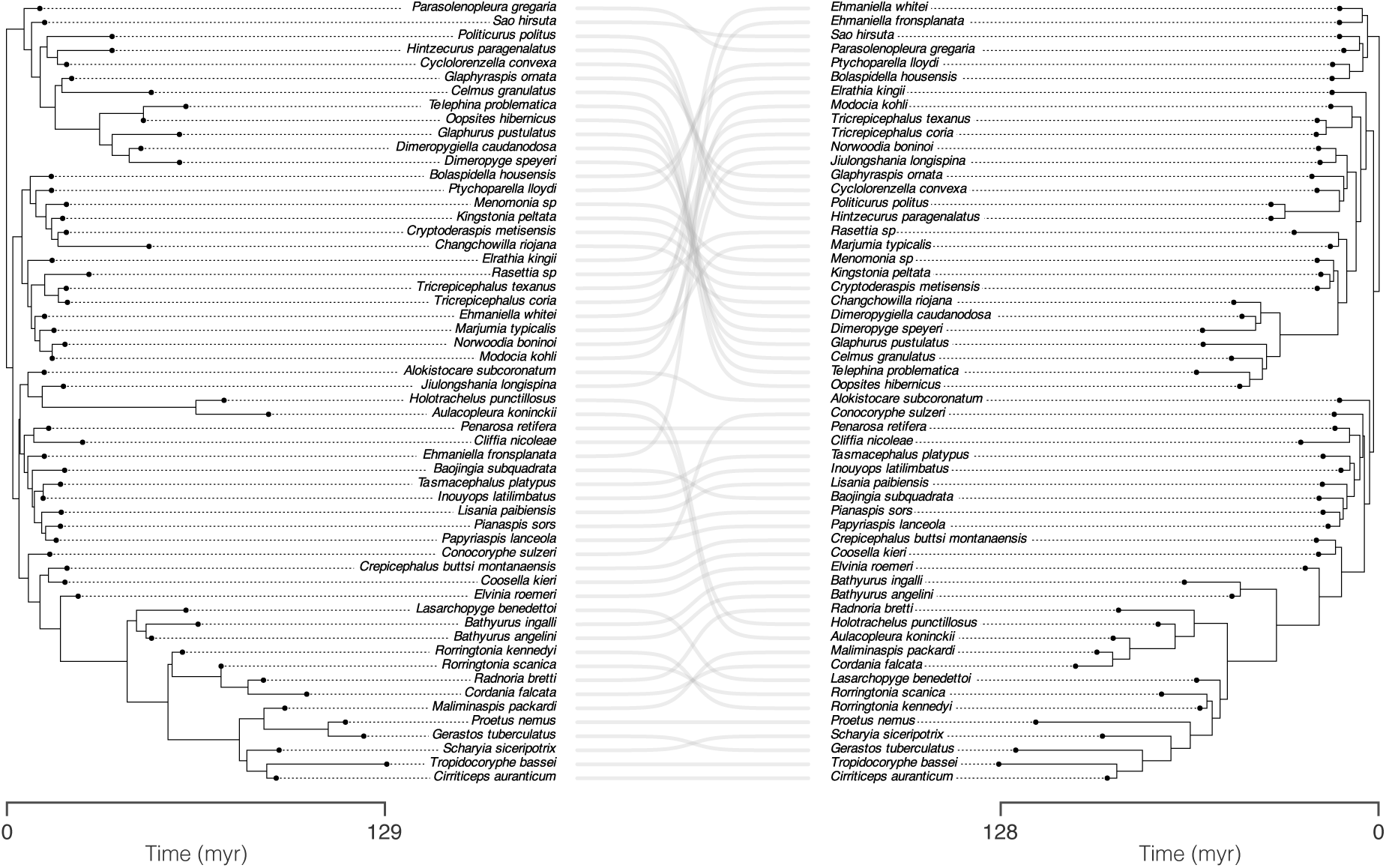
Cophylo plot of the maximum clade credibility trees from the resolved (left) and semi-resolved (right) showing the major topological (and consequently ghost range) differences.

## Notes

### Competing Interest Statement

The authors have declared no competing interest.

